# Diploidy confers genomic instability in *Schizosaccharomyces pombe*

**DOI:** 10.1101/2025.02.04.636513

**Authors:** Joshua M. Park, Daniel F. Pinski, Susan L. Forsburg

## Abstract

Whole genome duplication, or polyploidy, has been implicated in driving genome instability and tumorigenesis. Recent studies suggest that polyploidy in tumors promotes cancer genome evolution, progression, and chemoresistance resulting in worse prognosis of survival. The mechanisms by which whole genome duplications confer genome instability are not yet fully understood. In this study, we use *Schizosaccharomyces pombe* (fission yeast) diploids to investigate how whole genome duplication affects genome maintenance and response to stress. We find that *S. pombe* diploids are sensitive to replication stress and DNA damage, exhibit high levels of loss of heterozygosity, and become dependent on a group of ploidy-specific lethal genes for viability. These findings are observed in other eukaryotic models suggesting conserved consequences of polyploidy. We further investigate ploidy-specific lethal genes by depleting them using an auxin-inducible degron system to elucidate the mechanisms of genome maintenance in diploids. Overall, this work provides new insights on how whole genome duplications lead to genome instability.

## Introduction

Recent studies have estimated that ∼36% of tumors undergo a whole genome duplication event (Zack *et al*. 2013; Quinton *et al*. 2021). Whole genome duplications (polyploidy) have been increasingly associated with genome instability and tumorigenesis in various eukaryotic models (Fujiwara, *et al*. 2005; Storchová *et al*. 2006; Gemble *et al*. 2022). Human tetraploid cells show signs of increased DNA damage and chromosomal instability (Kuznetsova *et al*. 2015; Nano *et al*. 2019) that may promote cancer progression (Dewhurst *et al*. 2014). Polyploidy has also been linked to increased resistance of cancer cells to chemotherapeutic drugs (Mirzayans *et al*. 2018; Bharadwaj and Mandal 2020) and worse prognosis of survival in cancer patients (Dewhurst *et al*. 2014; Bielski *et al*. 2018).

Another hallmark of cancer is loss of heterozygosity (LOH; Steele *et al*. 2022). LOH is a phenomenon where a heterozygous region of the genome becomes homozygous, usually as a result of mitotic recombination (reviewed in Heil 2023). Crossover events or break-induced replication can result in long tracts of the genome losing heterozygosity, driving tumorigenesis through loss of tumor suppressor function (Had ija *et al*. 2001; Boulay *et al*. 2009; Lourenço *et al*. 2014; Ciani *et al*. 2022). Interestingly, cancer vulnerabilities can arise when non-driver genes undergo LOH, providing potential therapeutic targets (Nichols *et al*. 2020; Zhang and Sjöblom 2021). Recent research has suggested that cancer cells will undergo WGD as a means to combat the negative effects of LOH while still driving tumorigenesis (López *et al*. 2020; Archetti 2022). Although WGDs have been shown to play a role in genome instability and cancer progression, the underlying mechanisms require further investigation.

Yeasts provide a single celled model to study genome duplication. The budding yeast *S. cerevisiae* normally alternates between haploid and diploid states (Knop 2006), although higher degrees of polyploidy have been isolated and are associated with genome instability and LOH (Mayer and Aguilera 1990; Storchová *et al*. 2006; Storchová 2014; Sui *et al*. 2020; Dutta *et al*. 2022). In contrast, fission yeast is typically haploid, with a transient diploid zygote phase (Forsburg and Nurse 1991). In this organism, both diploids and higher level polyploids have been isolated but are unstable (Molnar and Sipiczki 1993).

In this study, we use the fission yeast to investigate the consequences of WGD, and the response to genomic stress. We compare *S. pombe* haploids to diploids and find that diploids exhibit increased sensitivity to genotoxins and experience elevated levels of LOH in the absence of exogenous stress. We show that this effect is independent of mating type heterozygosity. We also identify ploidy-specific lethal (PSL) genes that are conserved in budding yeast and humans (Storchová *et al*. 2006; Quinton *et al*. 2021). Using an auxin-inducible degron system, we investigate the mechanisms of cell death in PSL depleted diploids. We find that *S. pombe* diploids exhibit similar genomic instability phenotypes as human cells and tumors that have undergone WGD.

## Materials and Methods

### Yeast strains and growth

*S. pombe* strains used are listed in Table S1. *S. pombe* cells were cultured following standard protocols and methods (Forsburg and Rhind 2006).

### AID tagged strain construction

Auxin-inducible degron (AID) strains were constructed as described in Watson *et al*. 2021. pAW8-aid-5myc-Turg1-kanMX6 plasmid (Addgene 169356) was used as a template for PCR. Two rounds of PCR were performed to add sequence to both ends of the AID cassette that were homologous to the genomic target locus for C-terminus tagging. This PCR product was transformed using electroporation (Sabatinos and Forsburg 2010) into a wild type strain, then crossed with a strain with *OsTIR1F74A* integrated into the genome. Integration was validated by genomic PCR and Sanger sequencing. Rad52-AID-V5 strains were constructed by crosses using a strain provided by Dr. Antony Carr (Watson *et al*. 2021).

### Diploid strain generation

Mating type heterozygous diploid strains (sporulating and non-sporulating) were constructed using *ade6-M210/ade6-M216* intragenic complementation (Ekwall and Thon 2017). Mating type homozygous diploid strains were constructed by protoplast fusion (Park and Forsburg 2024). Diploid strains were validated by flow cytometery on the BD Accuri C6 Plus platform.

### Serial dilution growth assay

Liquid cultures were grown overnight in yeast extract +supplements (YES) media. Cultures were diluted to equal concentrations, then diluted five-fold serially. Dilutions were spotted onto YES or minimal glutamate media (PMG) plates with indicated supplements and drug concentrations. Plates were incubated at 32⁰C for the time indicated in figure legends.

### Acute genotoxic stress treatment survival

Liquid cultures were grown to mid log phase. Each culture was split in half for no treatment and treatment group. For irradiation (IR), cells were irradiated at the indicated doses using a 160 kV X-ray source (X-RAD iR-160, Precision X-Ray). For methyl methanesulfonate (MMS) treatment, MMS was added to the indicated doses and liquid cultures were incubated at 32⁰C with shaking for 4 hours. After treatment, cells were washed twice and resuspended in water. Cells were then counted on a Bright-Line Hemacytometer (Hausser Scientific) and 500 cells were plated onto YES + phloxine B plates. Plates were incubated at 32⁰C for 3 days, then colonies were counted and ratioed to untreated samples. Statistical significance was calculated using Mann-Whitney U-test (p-value > 0.05 ns, p-value < 0.05 *, p-value < 0.01 **, p-value < 0.001 ***, p-value < 0.0001 ****).

### Live cell fluorescent imaging

Fluorescent imaging was performed as described in Jones and Forsburg 2021. Briefly, liquid cultures were grown to mid-log phase in PMG supplemented with histidine, uracil, leucine, adenine, lysine, and arginine at 225 mg/L each (PMG-HULALA). MMS (0.0075%) or 5-Adamantyl-IAA (5’a-IAA, 100 nM) were added to the culture and incubated at 32⁰C with shaking. Samples were taken at the indicated timepoints for fluorescent imaging. Cells concentrated by a brief microfuge spin were applied to 2% agarose pads made from PMG + HULA and prepared on glass slides (Green *et al*. 2009). Images were acquired on a DeltaVision microscope with softWoRx v4.1 (GE, Issaquah, WA) using a 60x (NA1.4 PlanApo) lens, solid-state illuminator and 12-bit CCD camera. Images were acquired in twenty 0.2µm z-sections, then deconvolved and Maximum Intensity Projected (softWoRx, default settings). Images for publication were contrast adjusted using an equivalent histogram stretch on all samples. Color balance was adjusted and scale bars were added in Fiji (ImageJ 2.0). Statistical significance was calculated using Mann-Whitney U-test.

### Fixed cell imaging

Fixed cell imaging by 4′,6-diamidino-2-phenylindole (DAPI) staining was performed as described in Luche and Forsburg 2009. Cells were cultured in PMG-HULALA and samples were collected at the indicated time points. Samples were ethanol fixed in ice-cold 70% ethanol and rehydrated in water. 5 µL of rehydrated sample was placed on a microscope slide and dried on a hot plate (70⁰C). 5 µL of mounting solution (50% glycerol, 1 mg/mL p-Phenylenediamine, 1 µg/mL DAPI) was added on top of the dried sample and a cover slip placed on top. Images were acquired as described above.

### Induced mutagenesis and LOH assay

Liquid cultures were grown to log phase in YES media. Mutagenesis was induced by 0.0025% MMS treatment for one hour at 32⁰C with shaking. Cultures were unperturbed for the LOH assay. Cells were washed twice in water then counted on a hemacytometer. 500 cells were plated onto YES + phloxine B plates and 1×10^5^ cells were plated onto PMG supplemented with histidine and leucine (225 mg/L each), uracil (50 mg/L), and 5-Fluoroorotic acid (5-FOA, 1 mg/mL) plates (150 mm x 15 mm). YES + phloxine B plates were incubated for 3 days and 5-FOA plates for 4 days at 32⁰C before counting colonies. Haploidization was measured by counting lighter pink colonies on the phloxine B plates. 5-FOA resistant colonies were ratioed to YES colonies minus those that had haploidized. 5-FOA plates were replica plated onto YES plates containing nourseothricin or zeocin and incubated at 32⁰C overnight. Resistant colonies were ratioed to total 5-FOA resistant colonies. Statistical significance was calculated using Mann-Whitney U-test.

### LOH sequencing analysis

Hybrid diploid strains were constructed by mating a *mat2-102 S. pombe* strain containing a *ura4+* cassette with a *h-S. pombe var. kambucha* strain. Colonies that had undergone LOH were selected for using 5-FOA as detailed above. A control colony and 10 5-FOA resistant colonies were cultured in PMG supplemented with histidine, leucine, adenine, and uracil (225 mg/L each). 20 mL of mid log phase culture was processed for genomic DNA extraction using a Monarch HMW DNA Extraction Kit for Tissue (New England Biolabs). Library preparation was performed using a Native Barcoding Kit 24 V14 and samples were sequenced on a MinION Mk1C platform (Oxford Nanopore Technologies). Raw pod5 files were basecalled using Dorado v7.1.4 dna_r10.4.1_e8.2_400bps_5khz_sup model. Reads were aligned to the *S. pombe var. kambucha* reference genome and single nucleotide polymorphism (SNP) analysis was performed using the Geneious Prime 2024.0.7 software. SNP density plots were generated on R 4.3.1 with the package CMPlot (Yin *et al*. 2021).

### Protein extraction and immunoblot

Liquid cultures were grown to log phase, then treated with either 0.01% MMS (Sigma-Aldrich) or 10 nM 5’a-IAA (TCI America). Samples were collected at time points indicated and were washed twice in PBS. Sample pellets were then frozen in liquid nitrogen and all time point samples were processed together. Protein was extracted by trichloroacetic acid (TCA) precipitation (Grallert and Hagan 2017). Protein samples were quantified using Pierce BCA Protein Assay Kits (Thermo Scientific) and 25-50 µg protein was loaded. Samples were run on a 7.5% SDS-PAGE gel and transferred onto a nitrocellulose membrane. The following antibodies and dilutions were used for immunoblotting: α-*cdc2* (Novus Biologicals NB100-2716; 1:2000), α-V5 (Abcam ab27671; 1:1000), α-myc (Abcam ab9106 and Novus Biologicals NBP2-52636; 1:1000), α -Alpha Tubulin (Millipore Sigma T5168; 1:1000), α-Rabbit Alexa Fluor 488 (Thermo Scientific A32790; 1:1000), and α-Mouse Alexa Fluor 488 (Thermo Scientific A28175; 1:1000). Blots were visualized on the Amersham Typhoon platform and quantified using ImageQuant TL software (GE healthcare).

## Results

To study how diploidy affects genome stability in *S. pombe*, and to eliminate any contribution of mating type heterozygosity, we generated three kinds of diploids. First, we constructed a mating-type heterozygous *h^+^/h^-^* diploid that is sporulation competent. The second diploid h^-^ */mat2-102* lacks *matP*-information required for sporulation (Willer *et al*. 1995) but otherwise maintains mating type heterozygosity. The third strain is mating type homozygous, constructed by protoplast fusion. Initially, we examined *h^+^/ h^+^* homozygous strains, but to eliminate any potential reversion to *h^90^*, we also examined *h*^-^ *smt-0* / *h^-^ smt-0* strains that lack the site for the mating type switching imprint (Styrkársdóttir *et al*. 1993). These configurations control for any possible effects from mating type heterozygosity and isolate ploidy specific effects.

### Diploids are more sensitive to genotoxins

We observed that all three diploid strains exhibited increased sensitivity to a variety of genotoxic agents compared to the haploids (Figure 1A). All four diploid strains had similar levels of sensitivity, suggesting that mating heterozygosity did not have any affect, and *h^+^*/*h^+^* and *h^-^ smt-0*/*h^-^ smt-0* diploids behaved similarly (Figure 1A). We also observed increased sensitivity to irradiation in diploids compared to haploids across a range of exposure (Figure 1B). Because the *h^+^/h^-^*strain is prone to enter meiosis, we continued with the two non-sporulating strains for further characterization.

**Figure 1.**
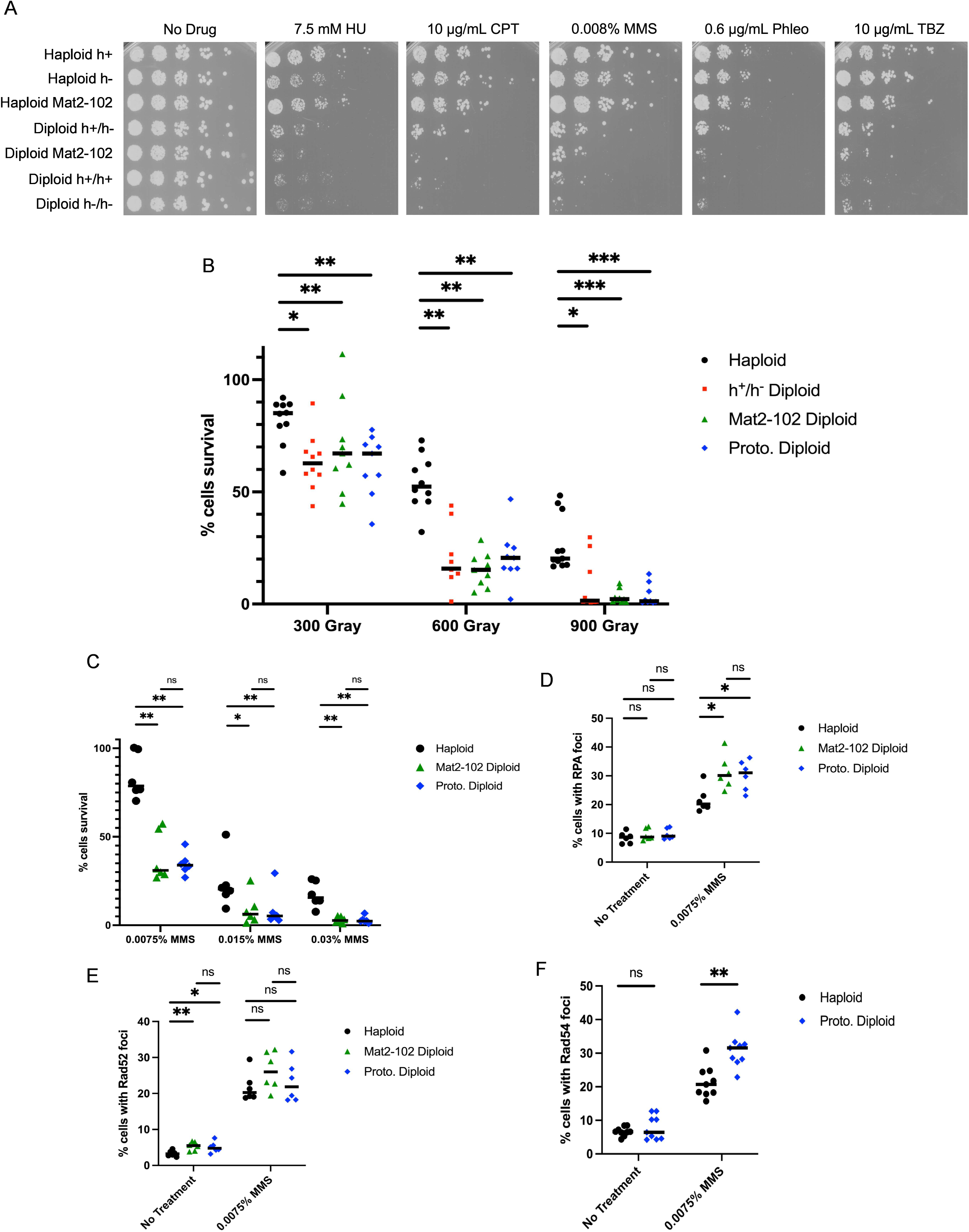
Diploids are sensitive to genotoxic stress. A) Serial dilution growth assay of haploid and diploid *S. pombe* strains. Diploid strains constructed by mating were crossed fresh before plating. Strains were spotted onto YES plates containing drugs as noted in the figure (HU=hydroxyurea, Phleo=phleomycin, CPT=camptothecin, MMS= methane methylsulfonate). Plates were incubated at 32⁰C for 3 days then imaged. B) Graph of haploid and diploid survival after irradiation. Statistical analysis was performed as mentioned in Materials and Methods. C) Graph of haploid and diploid survival after acute MMS treatment. *h^+^/h^-^* diploid was not included in this assay and future assays as there were no phenotypic differences with the Mat2-102 diploid, which is sufficient to control for mating type heterozygosity. D) Graph of RPA foci (*rad11-Cerulean fluorescent protein* [CFP]) after treatment with 0.0075% MMS, six replicates. E) Graph of Rad52 foci (*rad22-CFP*) after treatment with MMS, six replicates. F) Graph of Rad54 foci (*rad54-Green fluorescent protein* [GFP]) after treatment with MMS, nine replicates. Mat2-102 diploid was not included in this assay as there were no phenotypic differences with the protoplast fusion diploid.

We tested diploid response to acute treatment of the alkylating agent methyl methanesulfonate (MMS) and observed that diploids exhibited increased sensitivity across a range of concentrations (Figure 1C). We visualized DNA damage/repair foci using fluorescently tagged Ssb1 (RPA), Rad52, and Rad54 (Lisby *et al*. 2004; Sabatinos *et al*. 2012). We found that the percentage of cells with RPA and Rad54 foci did not significantly differ in haploid and diploid strains in unperturbed conditions (Figure 1D and 1F). After treatment with MMS, diploid cells showed an increased number of cells with RPA and Rad54 foci (Figure 1D and 1F). Interestingly, Rad52 foci were slightly elevated in diploids compared to haploids in unperturbed conditions but were at similar levels after MMS treatment (Figure 1E).

### Diploids have loss of heterozygosity

To further characterize the response of diploids to MMS, we compared the induced mutagenesis rate between haploids and diploids. We performed a forward mutation assay in *ura4+* strains by determining the frequency of 5-Fluoroorotic acid (5-FOA) resistance with and without MMS treatment (Liu *et al*. 1999; Dolan *et al*. 2010). We constructed diploid strains that were homozygous or heterozygous for the *ura4+* gene at its endogenous locus and treated haploid and diploid strains with 0.0025% MMS for one hour. To our surprise, the *ura4+* heterozygous strains had significantly elevated levels of 5-FOA resistance in both the untreated and treated conditions (Fig. 2A). We confirmed by PCR that nearly all the diploid samples tested had lost the entire *ura4+* gene (Table S2), resulting in a loss of heterozygosity (LOH) rather than point mutation(s).

**Figure 2.**
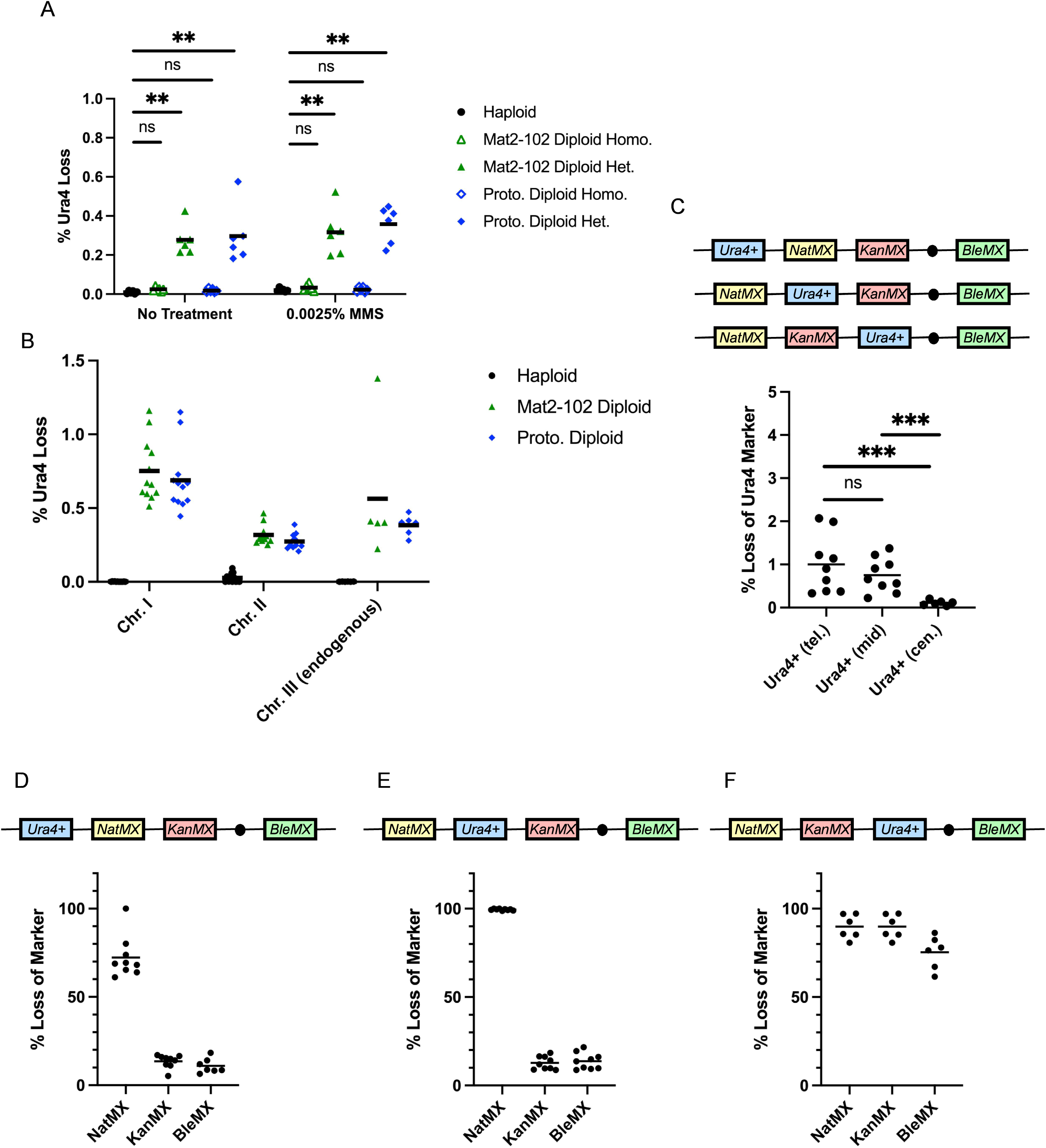
Diploids experience LOH in the absence of exogenous stress. A) Graph of induced mutagenesis in diploid strains that were heterozygous or homozygous for endogenous *ura4+* marker, six replicates. B) Graph of *ura4+* loss in diploid strains with *ura4+* inserted in chromosome I or II as heterozygous. *ura4+* was inserted ∼1.1 Mb downstream of the telomere of each chromosome compared to ∼150 kb downstream of the telomere on the endogenous chromosome III. C) Graph comparing *ura4+* loss when the marker was placed telomere proximal (tel.), middle (mid), or centromere proximal (cen.) on chromosome I, nine replicates. Middle marker was inserted 1.4 Mb downstream of the telomere proximal marker and the centromere proximal marker was inserted 1.2 Mb downstream of the middle marker. Marker on the right arm of chromosome I was inserted 800 kb downstream of the centromere. D-F) Graph showing loss of the other markers in the population that had lost the *ura4+* marker, nine replicates.

We were concerned that the endogenous *ura4+* locus’s proximity to the telomeres and rDNA array on chromosome III may have influenced the results, so we inserted the *ura4+* cassette into chromosome I and II. Again, we observed elevated levels of LOH in these diploids even in the absence of MMS (Fig. 2B). To assess whether this was a limited gene conversion, or loss of a chromosome, we constructed strains with the *ura4+* cassette and two drug markers *natMX* and *kanMX* (conferring resistance to nourseothricin and G418, respectively) equally spaced along the left arm of chromosome I and another marker *bleMX* (conferring resistance to bleomycin and related drugs) on the right arm. We observed that the rate of *ura4+* loss was similar in strains with the *ura4+* cassette inserted in the left and middle arm of the chromosome (Fig. 2C). However, the *ura4+* loss was significantly reduced in the strain that had the *ura4+* cassette inserted near the centromere (Fig. 2C).

Next, we measured the loss of the other markers in cells that had lost the *ura4+* cassette. In the strain with the *ura4+* telomere proximal, the middle marker was also lost in ∼70% of the population while the centromere proximal and right arm marker were lost in ∼10% (Fig. 2D). In the strain with the *ura4+* at the middle position, the telomere proximal maker was lost in ∼100% (Fig. 2E). However, in the limited population of colonies that had lost the centromere proximal *ura4+* cassette, ∼90% of the telomere proximal and middle markers were lost as well as ∼80% of the right arm marker (Fig. 2F). We suggest the high rates of LOH of the telomere-proximal and middle marker reflect an extended recombination or break induced replication event, while the loss of the centromere proximal *ura4+* reflects a chromosome mis-segregation event that results in homozygosis. Remarkably, this happens in strains that are otherwise unperturbed by genotoxic stress.

### Mapping LOH breakpoints in hybrid *S. pombe* diploids

The high rate of LOH associated with replication stress led us to wonder if there were recombination hotspots linked to fragile sites in the genome. To investigate recombination breakpoints, we used the *S. pombe variant kambucha* to generate a hybrid diploid with our *S. pombe* lab strain. The *kambucha* variant has less than 1% nucleotide divergence from *S. pombe* (Rhind *et al*. 2011) but provides enough single nucleotide polymorphisms (SNPs) to measure LOH in the hybrid diploid strain. We inserted the *ura4+* cassette in the left arm of Chromosome I in the *S. pombe* strain (Fig. 3A) and measured loss of the marker using 5-FOA as described above. We found that the hybrid diploid has slightly lower levels of LOH compared to an isogenic *S. pombe* diploid (Fig. S1A). We sequenced a control colony and 10 5-FOA resistant colonies, aligned the reads to the *S. pombe var. kambucha* reference genome, and performed a SNP analysis. Loss of the *ura4+* cassette on the *S. pombe* strain should be the result of recombination which leads to homozygosis of a region of the *S. pombe* chromosome and loss of SNPs in reference to the *S. pombe var. kambucha* genome. Indeed, we find that the 5-FOA resistant colonies have significantly reduced number of SNPs on the left arm of chromosome I (Fig. 3B) resulting in loss of heterozygosity. There were two classes of LOH in the 10 samples we analyzed: 7 of the 10 samples experienced a distinct break at ∼1.1 Mb and the remaining 3 samples experienced more extensive LOH (>2 Mb; Fig. 3B). The 1.1 Mb breakpoint coincides with a long terminal repeat element (LTR; SPLTRA.22) and an early firing origin 22 kb upstream of the breakpoint (Fig. 3C, S1B).

**Figure 3.**
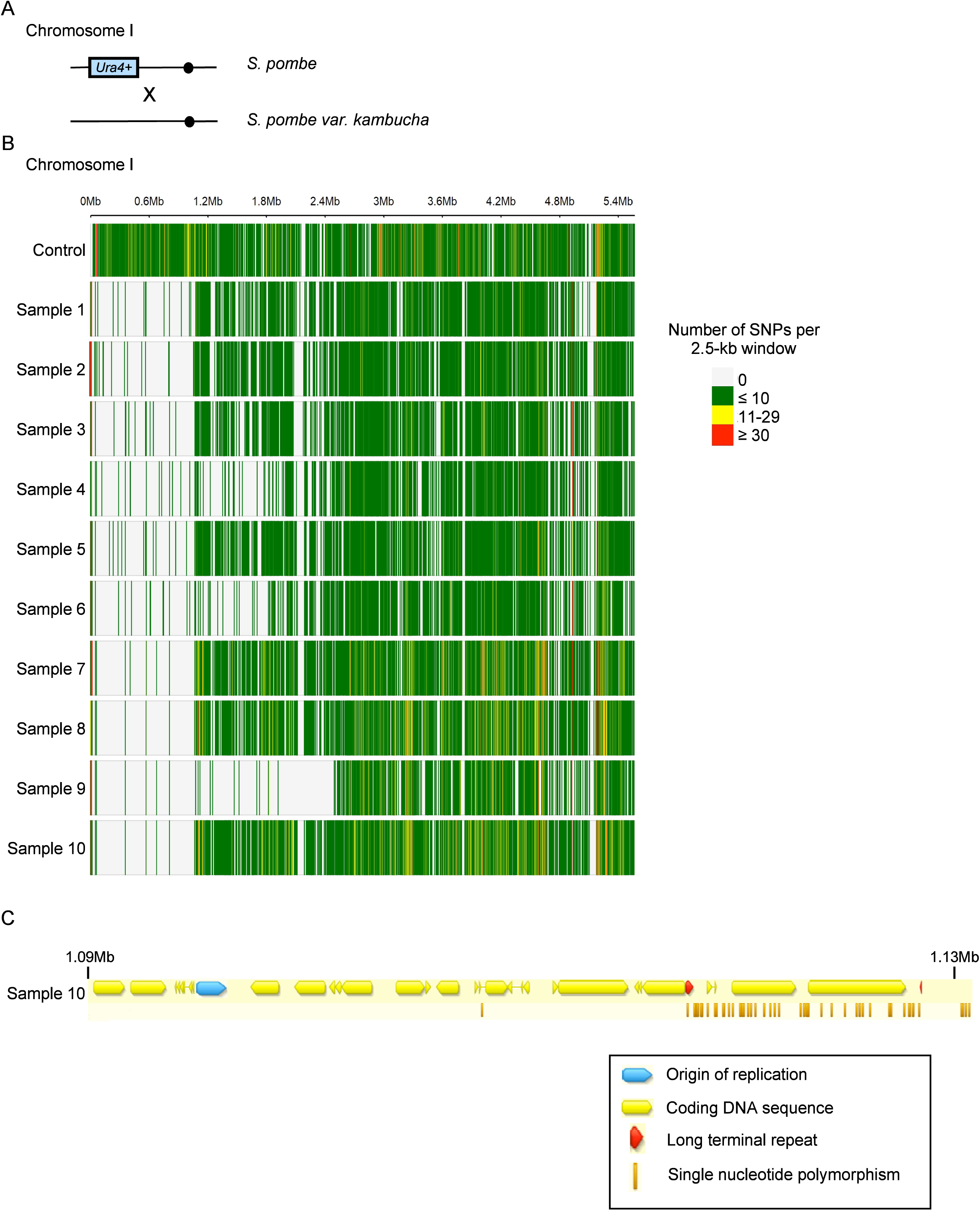
Sequencing of LOH samples show two distinct breakpoint sites A) Schematic of hybrid strain construction. B) SNP density plots depicting number of SNPs per 2.5 kb window across Chromosome I. C) Enlargement of Chromosome I at the 1.1 Mb breakpoint showing relevant gene features and SNPs of sample 10.

### LOH increases in mutant diploids

To investigate the pathways that may be involved with the LOH phenotype, we took a candidate approach. We selected a panel of strains with deletion mutations in genes that are implicated in genome maintenance pathways. We constructed homozygous deletion diploids with a single telomere proximal *ura4+* marker on the left chromosome arm, a marker in the middle of the left chromosome arm, and a marker on the right chromosome arm and assessed their effect on LOH frequency (Fig. 4A, Table 1). We found that deletion of checkpoint kinases *tel1Δ* and *chk1Δ* did not change the frequency of *ura4+* loss, but *rad3Δ* and *cds1Δ* diploids had a significant increase (1.4 and 3.2 fold increase, respectively). Deletion of *mrc1*, a replication checkpoint mediator and interactor of the fork protection complex (FPC; Tanaka and Russell 2001; Tanaka *et al*. 2010), also increased the loss of *ura4+* at a similar rate as *cds1Δ*. This suggests that an active replication checkpoint reduces the frequency of mitotic recombination in wild type cells.

**Figure 4.**
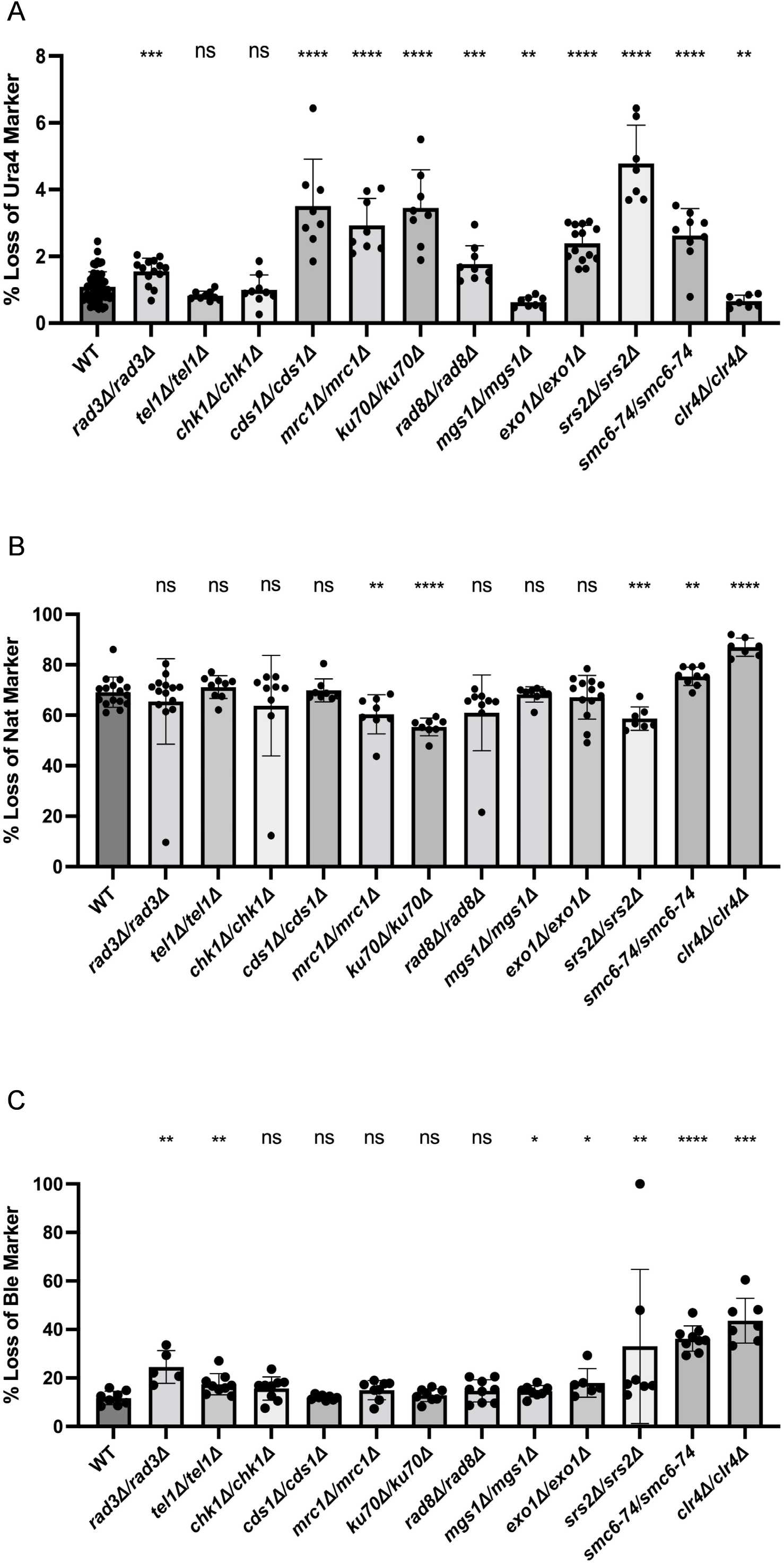
LOH in diploids is influenced by replication stress response and DNA repair genes. A) Bar graph showing rate of *ura4+* loss in homozygous deletion and mutant diploid strains. Bars represent mean and error bars represent standard deviation. Wild type (WT) had fifty four replicates. Mutant strains had seven to fourteen replicates due to picking colonies that already experienced LOH (both loss and homozygosing of the *ura4+* marker). B) Bar graph depicting rate of *natMX* loss (middle marker). C) Bar graph depicting rate of *bleMX* loss (right arm marker) indicating whole chromosome loss.

**Table 1.** List of genes tested for ploidy-specific lethality.

| Table 1 | List of genes tested for ploidy-specific lethality |  |
| --- | --- | --- |
|  | Gene | Function |
| *Diploid - Essential* |  |  |
|  | *Rad51* | Homologous recombination |
|  | *Rad52* | Homologous recombination |
|  | *Rad54* | Homologous recombination |
|  | *Mre11* | Resection |
|  | *Fbh1* | Helicase |
|  | *Mus81* | Structure specific endonuclease |
|  | *Crb2* | DNA end binding protein; checkpoint |
|  | *Top1* | Topoisomerase |
|  | *Swi3 (TIPIN)* | Fork protection complex |
|  | *Bub1* | Spindle assembly checkpoint |
|  | *Mad2* | Spindle assembly checkpoint |
|  | *Mad3* | Spindle assembly checkpoint |
| *Diploid - Unstable* |  |  |
|  | *Rqh1 (BLM)* | Helicase |
|  | *cdc27-D1* | Polymerase delta subunit required for BIR |
|  | *smc6-x* | DNA repair; checkpoint |
|  | *Swi1 (TIMELESS)* | Fork protection complex |
|  | *cdc2-asM17* | Cell cycle; |
|  | *cdc10-V50* | Cell cycle; MluI binding factor |
|  | *cdc25-22* | Cell cycle; phosphatase |
| *Diploid - Dispensable* |  |  |
|  | *Rad3 (ATR)* | Checkpoint |
|  | *Cds1* | Checkpoint |
|  | *Tel1 (ATM)* | Checkpoint |
|  | *Chk1* | Checkpoint |
|  | *Mrc1 (CLSPN)* | Fork protection complex |
|  | *Ku70* | Non-homologous end joining |
|  | *Lig4* | Non-homologous end joining |
|  | *Rad8* | HLTF helicase; ubiquitin ligase |
|  | *Rad16 (XPF)* | Structure specific endonuclease |
|  | *Mgs1 (WRNIP1)* | DNA-dependent ATPase |
|  | *Exo1* | Long range resection |
|  | *Srs2* | Helicase; Rad51 regulator |
|  | *smc6-74* | DNA repair; checkpoint |
|  | *Clr4* | Lysine methyltransferase |
|  | *Fml1 (FANCM)\** | Helicase |
|  | *Fml1/Fml2\** | Helicase |
|  | *top2-191\** | Topoisomerase |
|  | *Rev3\** | Translesion synthesis polymerase |
|  | *Eso1/Kpa1/Rev1/Rev3\** | Translesion synthesis polymerases |
|  | *Mus7\** | DNA repair |

List of genes that were tested for ploidy-specific lethality in *S.pombe* diploids. Diploid – essential genes were genes whose deletions or mutations were not able to be isolated as homozygous diploids by crosses or protoplast fusion. Diploid – unstable genes were genes whose deletions or mutations were able to be isolated as homozygous diploids, but failed to maintain diploidy over time (i.e. overnight culturing or freezing to add to collection resulted in haploidy). Diploid – dispensable genes were genes that were able to form stable diploids as homozygous deletions or mutations. Genes in the Diploid – dispensable category with asterisks (\*) were not included in LOH assay.

Next, we looked at a component of non-homologous end joining (NHEJ) *pku70*. We find that *pku70Δ* increases *ura4+* loss rate (3.2 fold increase), similar to *cds1Δ* and *mrc1Δ*. Deletion of *rad8*, an ubiquitin ligase (Frampton *et al*. 2006; Ding and Forsburg 2014), increased the rate of *ura4+* loss by 1.6 times compared to wild type (WT). Next, we looked at deletion of *mgs1*, a Werner helicase-interacting protein 1 (WRNIP1) homolog that is implicated in genome maintenance and DNA repair (Hishida *et al*. 2001; Saugar *et al*. 2012). Surprisingly, *mgs1Δ* resulted in a small but significant reduction in *ura4+* loss rate compared to the WT diploid (0.6 times lower).

We assayed HDR associated genes including *exo1Δ, srs2Δ, smc6-74.* Deletion of *exo1*, a long range resection exonuclease (Mimitou and Symington 2008), and *smc6-74*, a loss of function mutation in a structural maintenance of chromosome (SMC) gene involved in HDR (Ampatzidou *et al*. 2006; Irmisch *et al*. 2009), both increased the rate of loss of *ura4+* by 2.2 and 2.4 fold, respectively. Deletion of *srs2*, a helicase with anti-recombinogenic properties (Doe and Whitby 2004), induced the highest increase in *ura4+* loss among all the mutants tested at 4.4 fold. Finally, we looked at deletion of *clr4*, a lysine methyltransferase responsible for depositing histone H3K9 methylation at the pericentromeric heterochromatin (reviewed in Allshire and Ekwall 2015). Although we find that *clr4Δ* reduces loss of *ura4+*, this may be skewed by the high rate of haploidization in this strain probably reflecting chromosome loss due to centromere disruption (Fig. S2). Overall, we find a trend that genes involved in response to replication stress and DNA repair tend to increase the loss of the *ura4+* marker.

In the population of diploids that had lost the *ura4+* marker, we also probed for the loss of the other markers on the left and on the right arm of the chromosome. We observed that *mrc1Δ*, *pku70Δ*, and *srs2Δ* have reduced rate of loss of the additional markers compared to WT, while *smc6-74* and *clr4Δ* have increased rates (Fig. 4B). For the marker on the right arm of the chromosome, we found that *smc6-74* and *clr4Δ* had the largest increase in rate of loss (Fig. 4C). This is consistent with *clr4* and *smc6-74* impacting loss of the entire chromosome, which may reflect whole chromosome mis-segragation. Notably, *srs2Δ* had several outliers in both directions of rate change suggesting it may impact multiple pathways.

Next, we tested MMS sensitivity for haploid and diploids mutants used in the LOH assay. We found that *exo1Δ* and *smc6-74* diploids were more sensitive to MMS than wild type diploids and their haploid counterparts (Fig. S3). The remaining diploid mutants were either just as sensitive to MMS as their haploid counterparts, or comparable to the wild type.

### Mutants inviable as diploids

Previous studies suggested that deletions of DNA homologous recombination genes *rad51, rad52,* and *rad54* could not be constructed as diploids (Catlett and Forsburg 2003; Grishchuk *et al*. 2004; Octobre *et al*. 2008). Similar results were observed for *bub1* (Bernard *et al*. 1998). In the process of constructing diploid mutants for the LOH assay, we validated and identified new ploidy specific lethal genes in *S. pombe* (Table 1). We confirmed that *rad51Δ, rad52Δ,* and *rad54Δ* diploids could not be stably isolated by crossing or protoplast fusion. Additionally, we found *mre11Δ, fbh1Δ, mus81Δ,* and *crb2Δ* diploids could not be constructed, adding evidence to the homologous recombination pathway being essential in *S. pombe* diploids. We also found that deletion of topoisomerase I *top1* and FPC component *swi3* were inviable, suggesting that responding to replication stress is essential as ploidy increases. We observed that spindle assembly checkpoint genes *bub1, mad2,* and *mad3* are also essential in diploids. For certain deletions, we were initially able to isolate diploids under selection but were unable to maintain them as they haploidized in culture. High frequency of haploidization is linked to chromosome loss (Bodi *et al*. 1991). These included additional DNA repair and FPC genes (*rqh1Δ, smc6-x, cdc27-D1,* and *swi1Δ*) and cell cycle mutants generally used for synchrony (*cdc10-V50, cdc25-22,* and *cdc2-asM17*) (Table 1).

Seeing as these group of genes were essential for diploid viability, we were interested if overexpression of these genes could rescue diploid sensitivity to genome stress. We tested overexpression of the DNA damage and replication stress genes in response to MMS and the spindle assembly checkpoint genes in response to the microtubule destabilizing agent thiabendazole (TBZ). We found that overexpression of the diploid essential genes did not rescue diploid sensitivity to MMS or TBZ, and in some cases resulted in decreased viability in both haploids and diploids regardless of genotoxic stress (Fig. S4).

### Ploidy-specific lethal protein depletion in diploids

To investigate why some genes are necessary for survival in *S. pombe* diploids, we used the auxin-inducible degron (AID) system (Watson *et al*. 2021). Not all genes were amenable to this degron fusion but we successfully constructed diploid strains with both copies of *rad52* and *mad2* tagged with an AID cassette. In the haploid strains, the addition of the auxin analog 5’adamantyl-IAA (5’a-IAA) leads to rapid degradation of the tagged protein within 30 minutes (Watson *et al*. 2021, Fig. 5C). In the diploid strains, protein degradation was ablated for Mad2 but not Rad52 (Fig. 5A and 5C). However, both diploid strains showed significant decrease in viability when plated on media with 5’a-IAA (Fig. 5B and 5E) suggesting that the level of protein depletion was sufficient for loss of viability. The Rad52-AID diploids showed hypersensitivity to low levels of HU compared to their haploid counterpart when Rad52 was depleted (Fig. 5B). The AID cassette did increase sensitivity to MMS in Rad52 tagged haploid and diploid strains in the absence of 5’a-IAA, suggesting impairment of protein function (Fig. S5). Similarly, Mad2-AID diploids exhibited increased sensitivity to low levels of TBZ (Fig. 5E). These results highlight the effectiveness of using the AID system in *S. pombe* diploids and provides a tool for investigating the mechanisms on how depletion of PSL proteins leads to cell death.

**Figure 5.**
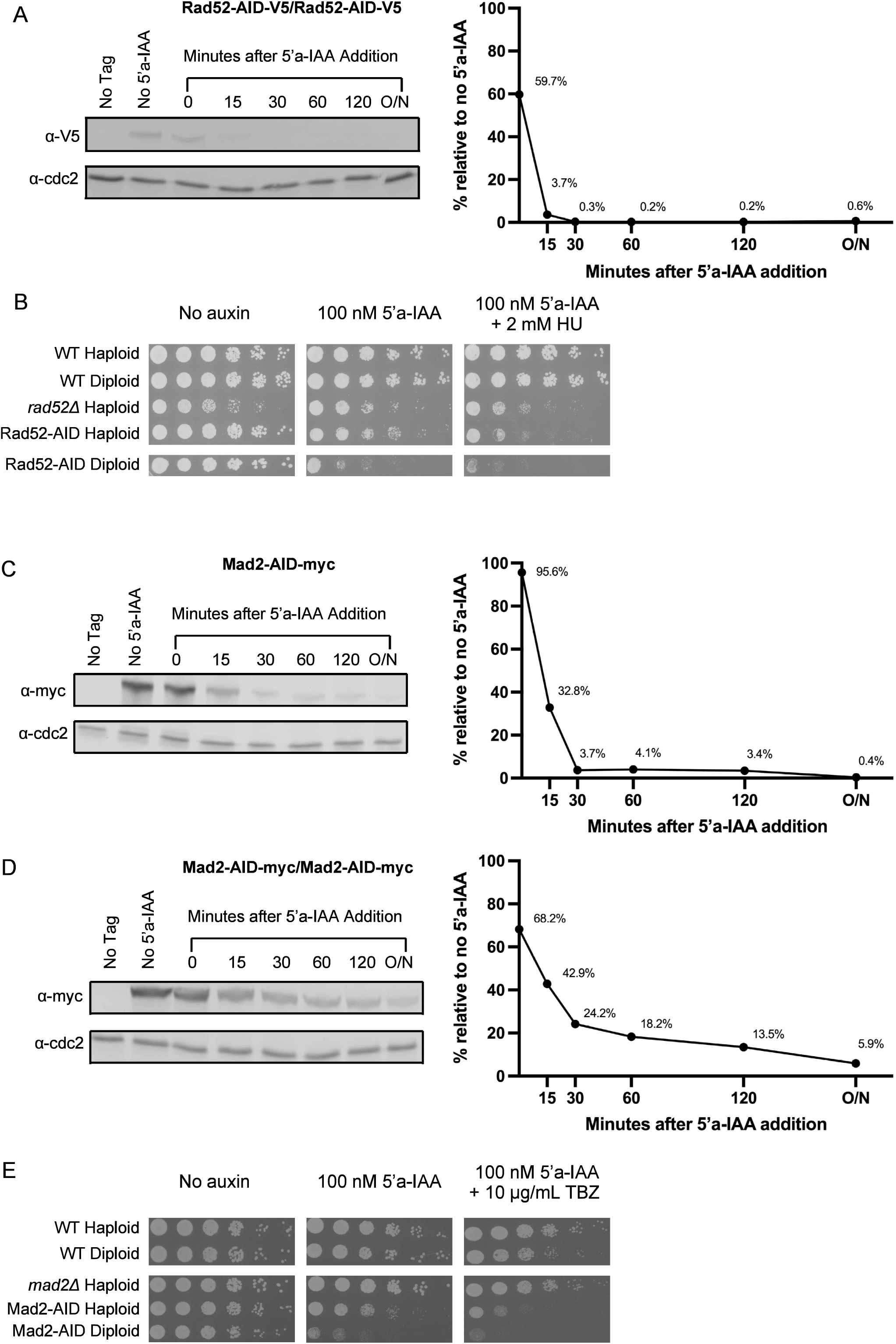
Depletion of Rad52 and Mad2 using the AID system results in loss of viability in diploids. A) Immunoblot and quantification of Rad52 degradation in Rad52-AID homozygous diploids. Cells were treated with 100 nM 5’a-IAA and sampled at the indicated time points. Cdc2 was used as the loading control. B) Control and Rad52-AID strains were spotted onto YES plates with and without 100 nM 5’a-IAA and/or 2 mM HU. Plates were incubated at 32⁰C for 3 days then imaged. C-D) Immunoblot and quantification of Mad2 degradation in Mad2-AID haploids and Mad2-AID homozygous diploids. Conditions were the same as the Rad52 assay. E) Control and Mad2-AID strains were spotted onto YES plates with and without 100 nM 5’a-IAA and/or 10 µg/mL thiabendazole (TBZ). Plates were incubated at 32⁰C for 3 days then imaged.

### Characterizing Rad52 depletion in diploids

We investigated the phenotype of Rad52 depletion, beginning with accumulation of RPA repair foci. Both haploids and diploids had increased number of cells with RPA foci at 4 hours of depletion, but the diploids had about 1.6 time more (39% and 65%, respectively; Fig. 6A). The number of cells with RPA foci steadily increased in both haploids and diploids with time but maintained a similar ratio. Diploids accumulated increasing levels of multi-foci RPA compared to haploids (Fig. S6A). While the haploids continued to divide and increase in cell number, diploids mainly grew in size (coinciding with increase in OD) and divided very slowly (Fig. 6B and 6C). We see that Chk1 is persistently activated when Rad52 is depleted in both haploid and diploid strains (Fig 6D and 6E). The Chk1 activation is strongest at the 4 hour mark, but is not as strong compared to an MMS treated sample.

**Figure 6.**
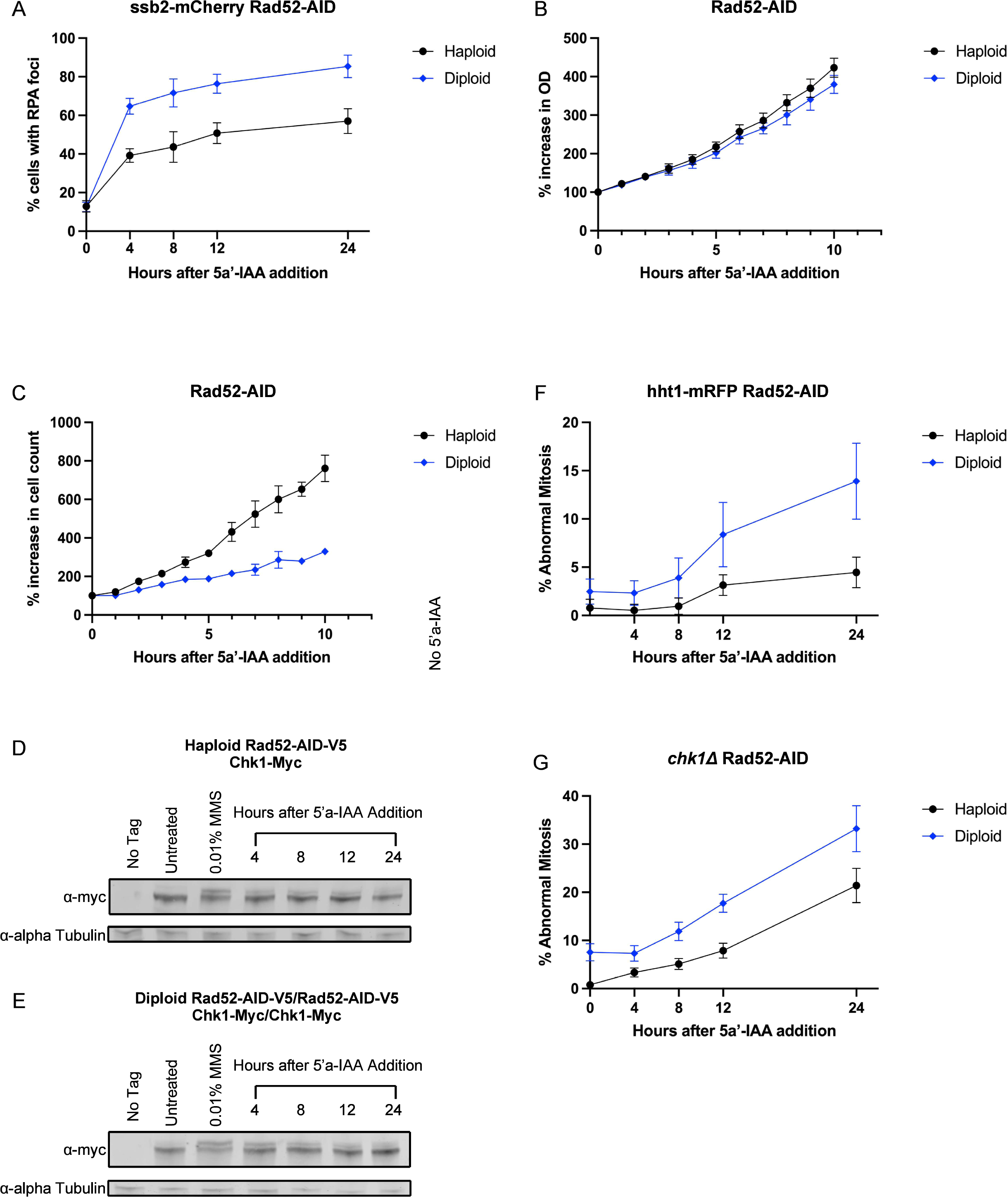
Depletion of Rad52 leads to increased genome instability in diploids. A) Graph of RPA foci (*ssb2-mCherry*) in haploid and diploid Rad52-AID strains after 5’a-IAA treatment. Samples were collected at the time points indicated and imaged by fluorescent microscopy, nine replicates per time point. B) Graph showing percent increase in optical density (OD; 595 nm) relative to the time point before 5’a-IAA addition. Time points were taken every hour for ten hours, three replicates. C) Graph showing percent increase in cell count relative to time point before 5’a-IAA addition. Samples were from the same batch used for OD reading. D-E) Immunoblot of Chk1 activation in Rad52-AID haploids and Rad52-AID homozygous diploids. Cells were treated with 100 nm 5’a-IAA and sampled at the indicated times. Alpha tubulin was used as the loading control. F) Graph showing percent increase in abnormal mitotic events measured using a histone marker tagged with a red fluorescent protein (*hht1-RFP*). Samples were collected at the time points indicated and imaged by fluorescent microscopy, nine replicates per time point. G) Graph showing percent increase in abnormal mitotic events measured using DAPI stain on fixed cell samples collected at the time points indicated. Nine replicates per time point.

We used a fluorescently labeled histone to measure any abnormal nuclear morphology and mitotic events (e.g. lagging chromosomes, uneven segregation, micronuclei; Fig. S6B and S6C). Most of the diploids showed single nuclei consistent with cell cycle arrest. Diploids had a small increase in nuclear abnormalities before treatment with 5a’-IAA, and the difference grew throughout the time course (Fig. 6F). We observed that at the 12 and 24 hour marks the diploids exhibited aberrant septation attempts such as multiple septa (Fig. S6D). These phenotypes are consistent with prolonged cell cycle arrest. Next, we deleted Chk1 in Rad52-AID strains and visualized DNA by DAPI staining in fixed cells as we were unable to isolate *chk1Δ* Rad52-AID histone tagged strains. We find a similar pattern of increasing levels of abnormal mitotic events as time progresses, but to a higher degree compared to the *Chk1^+^* Rad52-AID strains (Fig. 6G). Interestingly, *chk1Δ* Rad52-AID diploids do not arrest at the 12 hour mark like the *Chk1^+^* Rad52-AID diploids do but eventually seem to arrest at the 24 hour mark (Fig. S6E). These results suggest that DNA damage and Chk1 activation upon Rad52 depletion passes the threshold for cell cycle arrest in diploids but not haploids.

## Discussion

In this study, we investigate how diploidy affects genome stability in the fission yeast *Schizosaccharomyces pombe*. This organism is typically a haploid, and generally only forms diploids as brief zygotes before entering meiosis. However, vegetative diploids can be recovered in the laboratory using complementing markers. This is in contrast to budding yeast, which alternates between diploid and haploid states.

In budding yeast, mating type heterozygosity affects DNA repair in diploids (Kadyk and Hartwell 1992; Heude and Fabre 1993; Barbour and Xiao 2006). We compared *S. pombe* diploids with different configurations of mating type loci from fully heterozygous, to fully homozygous, and observed no difference in growth or sensitivity to genotoxins. All *S. pombe* diploids exhibit increased sensitivity to a wide range of genotoxins regardless of mating type heterozygosity (Fig. 1A). This result was unexpected as we hypothesized that having another set of chromosomes would provide an additional template for homology directed repair and act as a buffer for recessive mutations that would be deleterious in a haploid context. Early work in *S. cerevisiae* showed that diploids exhibited increased resistance to irradiation compared to haploids, but further increasing ploidy resulted in decreased resistance (Mortimer 1958). However, more recent work in budding yeast and humans found that tetraploids have increased sensitivity to replication stressors and double-strand break inducing agents (Storchová *et al*. 2006; Wangsa *et al*. 2018). Our results suggest that increase in ploidy from the natural genome state confers genomic instability in *S. pombe* as well, although in this case it is from haploid to diploid, rather than diploids to tetraploid as in budding yeast and humans.

We examined diploid response to the genotoxin methyl methanesulfonate (MMS). MMS is an alkylating agent that creates DNA adducts which leads to replication stalling (Szyjka *et al*. 2008; Iyer and Rhind 2017) and induces a replication stress transcriptional response in both haploids and diploids (Park and Forsburg 2024). However, the degree of transcriptional induction of damage responsive genes is somewhat attenuated in diploids (Park and Forsburg 2024). We found that an acute treatment of MMS reduces viability in diploids (Fig. 1C), consistent with an impaired damage response. When we look at DNA damage markers, we find that diploids have a slight increase in cells with Rad52 foci compared to haploids (Fig. 1E) but not RPA or Rad54 (Fig. 1D and 1F) in unperturbed conditions. This is in contrast with *S. cerevisiae* and human tetraploids that have increased DNA damage markers (Storchová *et al*. 2006; Gemble *et al*. 2022). Upon treatment of MMS, *S. pombe* diploids have increased DNA damage markers compared to untreated diploids (Fig. 1D-F). These results, taken together with the increased sensitivity of diploids to MMS, are consistent with *S. pombe* diploids having an impaired repair response and increased damage in MMS.

When yeast cells are treated with low levels of MMS, the cells will repair some of the lesions using an error-prone pathway involving translesion polymerases that can be measured using a forward mutation assay (Stelter and Ulrich 2003; Dolan *et al*. 2010). We observed no change in induced mutagenesis in diploids, but to our surprise, we found that our diploid strains exhibited high levels of loss of *ura4+* function even in untreated cells (Fig. 2A). We determined the cause to be partial loss of heterozygosity (Table S2). The LOH phenomenon was observed for all three *S. pombe* chromosomes (Fig. 2B) and occurred at similar rates when the markers were inserted telomere proximal and in the middle of the chromosome arm (Fig. 2C) suggesting a global effect. *S. cerevisiae* diploids have also been shown to experience spontaneous LOH using similar systems, but at rates that are 2-4 magnitudes lower than our results (Chen *et al*. 1999; Ohnishi *et al*. 2004; Storchová *et al*. 2006). LOH in *S. cerevisiae* diploids were a result of crossover and break-induced replication, non-crossover gene conversion events, and chromosome loss (Ohnishi *et al*. 2004; Suetomi *et al*. 2010; Chumki *et al*. 2016). The majority of the LOH events in our assay resulted in the loss of the telomere proximal marker but maintained the opposite arm of the chromosome, with only a small fraction of events reflecting whole chromosome mis-segregation (Fig. 2E). A recent study in diploid fission yeast showed interactions between homologous chromosomes after an induced break that was Rad51 dependent (Vines *et al*. 2022). This suggests that HDR uses homologous chromosomes, in addition to sister chromatids, for repair in fission yeast. We infer that the LOH events reflect recombination or BIR and speculate that these may be driven by fragile sites that are prone to replication stress.

We created *S. pombe*/*S. pombe var. kambucha* diploids, in which the chromosomes can be distinguished by SNPs. We find there are two classes of LOH: a clear breakpoint at ∼1.1 Mb position and a more diffuse site closer to the centromere (Fig. 3B). The ∼1.1 Mb breakpoint coincides with a long terminal repeat in 7 of the 10 samples which may suggest a potential fragile site that is prone to recombination (Fig. 3C, S1B). This may be similar in nature to the known fragile site FS2 in *S. cerevisiae* associated with Ty1 elements that causes mitotic recombination in diploid budding yeast (Rosen *et al*. 2013). The other three samples exhibited much longer tracts of LOH, but did not share similar breakpoints between them. All three samples did have LTR elements in the vicinity of the breakpoint. These potential fragile sites may play a role in the increased endogenous replication stress that require HDR protein for repair. However, further investigation is required to determine if these LTR elements are fragile sites and if there are other sites in the *S. pombe* genome.

We see LOH rates increase in strains that lack replication checkpoint responders such as Mrc1 or Cds1. Deletion or mutation of genes involved in DNA repair (*rad8Δ, rad16Δ, exo1Δ, srs2Δ, smc6-74*) also increased LOH to varying degrees. Rad8 has been shown in *S. pombe* to play a role in responding to stalled replication forks through post replication repair (PRR) pathways (Frampton *et al*. 2006; Ding and Forsburg 2014). Knock out of Rad8 potentially drives repair through HDR, resulting in the increase in LOH observed. Similarly, deletion of Srs2 and Exo1 have increased rates of mitotic recombination in *S. cerevisiae* and *S. pombe* resulting in LOH (Wang *et al*. 2001; Ira *et al*. 2003; Lydeard *et al*. 2010; Marrero and Symington 2010). Interestingly, *smc6-x* mutants were inviable diploids, but using the *smc6-74* allele, we could construct diploids and observed elevated levels of LOH. The Smc5/6 complex has been implicated in processing stalled replication forks (Morikawa *et al*. 2004; Pebernard *et al*. 2006) and the difference in the two mutants may reflect a separation of function (Harvey *et al*. 2004) that is emphasized in diploids. Deletion of the NHEJ protein Ku70 also resulted in increased LOH. The ablation of NHEJ may drive increased repair by HDR resulting in mitotic recombination and LOH (Maruyama *et al*. 2015; Li *et al*. 2018). *mgs1Δ* resulted in a slight decrease of *ura4+* loss in our assay. Deletion of *MGS1* in *S. cerevisiae* haploids and diploids caused an increase in recombination (Hishida *et al*. 2001) in contrast to our results. Studies on Mgs1 in *S. pombe* are lacking and further investigation is required to elucidate the differences observed between budding and fission yeast. We have generally found that genes involved in responding to replication stress are either essential or lead to higher levels of LOH when deleted. This suggests the *S. pombe* diploids experience increased endogenous replication stress that lead to mitotic recombination and cell death if core HDR pathways are compromised.

As we were constructing the mutants for the LOH assay, we found that a subset of mutants could be isolated but not maintained as diploids due to rapid haploidization which typically occurs following whole chromosome loss (Bodi *et al*. 1991). These include a BIR-specific allele of DNA polymerase delta (*cdc27-D1*; Tinline-Purvis *et al*. 2009), a helicase that dissolves HR intermediates (*rqh1Δ*; Wu *et al*. 2006; Hope *et al*. 2007), and a member of the fork protection complex (*swi1Δ*). The histone methyltransferase Clr4, which is required for efficient chromosome segregation (Ekwall *et al*. 1996), also experienced high levels of haploidization but maintained enough diploids for the experimental assay (Fig. S2). Interestingly, cell cycle mutants *cdc10-V50, cdc25-22,* and *cdc2-asM17* could not be maintained as diploids even at permissive temperatures, suggesting *S. pombe* diploids are very sensitive to cell cycle perturbations.

We did not find any mutants that dramatically reduced the rate of LOH. We suggest this is related to the requirement of some genes for viability in diploids. The Rad52 epistasis group required for homologous recombination, as well as spindle assembly checkpoint mutants, have previously been reported to be inviable as diploids in fission yeast (Bernard *et al*. 1998; Catlett and Forsburg 2003; Grishchuk *et al*. 2004; Octobre *et al*. 2008). Interestingly, screens for ploidy-specific lethal genes in both budding yeast and human tetraploid cells have also identified HR components, kinetochore, and SAC genes (Storchová *et al*. 2006; Quinton *et al*. 2021). These studies suggested that kinetochore functions are particularly important in tetraploids and may not scale effectively, but the role of the HR proteins has not been investigated. We speculate that the LOH event is an essential process requiring HR proteins.

Using a candidate approach, we found that, in addition to the previously identified genes, additional fission yeast PSL mutants include topoisomerase I (*top1Δ*), the structure specific endonuclease *mu81Δ,* the helicase *fbh1Δ,* and the fork protection component *swi3*Δ. These are all involved in resolving replication fork collapse (Shyian *et al*. 2020; Grabarczyk 2022; reviewed in Chakraborty *et al*. 2023) and again suggest that diploid fission yeast have increased susceptibility to replication stress. We propose that the LOH reflects diploids-specific fragile regions on the chromosome.

Using an auxin-degron fusion to Rad52, we find that depletion of Rad52 leads to loss of viability, with increased RPA foci and elongated cells, most with a single nucleus, consistent with cell cycle arrest (Fig. 5B, 6, and S6). Although the haploids also exhibited persistent RPA foci, they did not show any signs of cell cycle arrest and had a minor increase in abnormal mitosis. These results suggest that haploids are capable of responding to the increased damage without inducing arrest when Rad52 is depleted, but the diploids pass the threshold of DNA damage that is manageable resulting in cell death.

Our previous transcriptome analysis suggested that most transcripts scale up ∼2x in diploids relative to haploids, but there may be modestly reduced expression of a few genes including the polymerase subunit *cdc1* (Park and Forsburg 2024). It is possible that the transcription doesn’t lead to an appropriately scaled response of all replication factors. Overexpression of individual diploid essential genes in fission yeast did not rescue MMS sensitivity (Fig. S4), suggesting a more global increase is required to see a response. A screen in fission yeast diploids found that heterozygous deletion of some replication factors, including *cdc1*, resulted in haploinsufficient growth (Kim *et al*. 2010). Indeed, a recent report from human cells suggests that the first cell cycle following tetraploidization is extremely prone to replication stress, due to insufficient dosage of some replication factors (Gemble *et al*. 2022). We speculate that fission yeast diploids may model this state.

Taken together, our data suggest that fission yeast diploidy has the same deleterious effects observed in budding yeast or human tetraploidy (Storchová *et al*. 2006; Quinton *et al*. 2021; Gemble *et al*. 2022). This suggests there is a conserved response to whole genome duplication above normal ploidy. Since diploids are genetically more straightforward in analysis, this establishes *S. pombe* as a useful model for polyploidy.

### Data Availability

The sequencing data discussed in this publication have been deposited in NCBI’s Sequence Read Archive under BioProject ID PRJNA1216252. Strain list is available in Table S1. All strains and reagents are available upon request.

## Acknowledgements

We thank Jiping Yuan and Kassandra Martinez for their technical assistance and members of the lab for helpful comments. We thank Dr. Nicholas Rhind and Dr. Sarah Zanders for their help and generosity with strains and resources for the *S. pombe var. kambucha* project. We thank Dr. Antony for providing strains for the auxin-inducible degron project. We thank Dr. Irene Chiolo was allowing us to use the X-RAD iR-160 machine.

## Competing interests

The authors declare no competing or financial interests.

## Funding

This research was supported by the National Institute of General Medical Sciences [R35-GM118109] (S.L.F) and University of Southern California Student Opportunities for Academic Research (D.F.P).

---

Figure S1. LOH rate of *S. pombe/S. pombe var. kambucha* hybrid strain

A) Graph showing the *Ura4+* loss rate in isogenic *S. pombe* diploids and hybrid *S. pombe/S. pombe var. kambucha* diploids. B) Enlargement of Chromosome I at the breakpoints of samples 1-9 showing relevant gene features and SNPs.

Figure S2. Haploidization rates of mutant diploid strains.

Bar graph showing haploidization rates in the mutant diploid strains used in the LOH assay. Haploids were measured as lighter pink colonies compared to the darker pink colonies when plated on phloxine B plates. Lighter colonies were ratioed to total number of colonies counted.

Figure S3. MMS sensitivity assay of the LOH mutant panel.

Serial dilution growth assay of mutant haploid and diploid strains used in the LOH assay. Strains were spotted onto pombe minimal glutamate (PMG) media supplemented with histidine, uracil, leucine, and adenine without MMS or with MMS at the concentrations as noted in the figure.

Plates were incubated at 32⁰C for 3 days then imaged.

Figure S4. Overexpression of diploid essential genes does not rescue MMS sensitivity

Serial dilution growth assay of haploid and diploid strains overexpressing diploid essential genes listed in Table 1. Yeast strains were transformed with a “no message in thiamine” (NMT) promoter vector controlling the expression of the gene of interest or an empty vector (EV) as control. Transformed yeast strains were spotted onto PMG media supplemented with histidine and leucine to maintain the vector (uracil selection) and ploidy (adenine selection). Thiamine was added to a final concentration of 5 µg/mL to repress expression or omitted to induce expression of the gene of interest. MMS was added at the concentrations as noted in the figure. Plates were incubated at 32⁰C for 3 days then imaged.

Figure S5. Rad52-AID strains exhibit MMS sensitivity in the absence of degradation.

Serial dilution growth assay of Rad52-AID haploid and diploids. Strains were spotted onto YES containing MMS at the indicated concentrations without 5’a-IAA. Plates were incubated at 32⁰C for 3 days then imaged.

Figure S6. Diploids exhibit abnormal nuclear and cell phenotypes when Rad52 is depleted.

A) Stacked bar graph showing distribution of single, multiple, and large RPA (*ssb2-mCherry*) foci ratios in haploid and diploid Rad52-AID strains after 5’a-IAA addition. Two or more RPA foci were categorized as multiple foci and RPA foci that were amorphous and larger on average were categorized as large foci. B) Representative image of diploid cell with RPA coated bridge at eight hours after 5’a-IAA addition. C) Representative image of diploid cells with abnormal segregation of DNA twelve hours after 5’a-IAA addition. D) Representative brightfield image of diploid cells with abnormal septation patterns twenty four hours after 5’a-IAA addition. E) Representative brightfield images of *Chk1^+^* and *chk1Δ* Rad52-AID cells at the time points indicated.

**Table S2.**
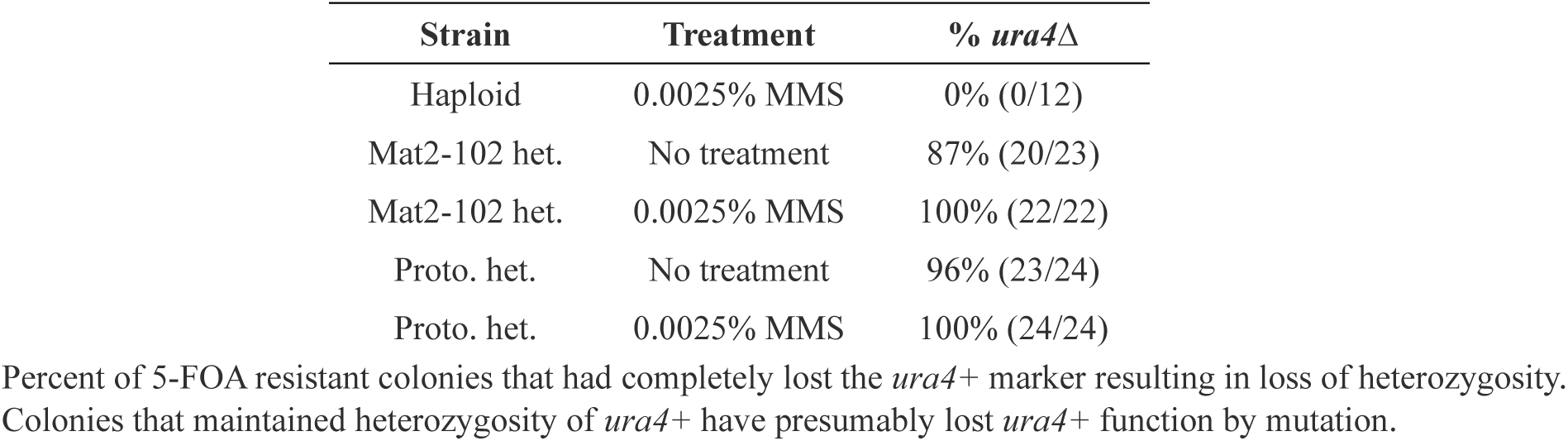
Percent loss of *ura4+* in diploids.

| Table S2 |  | Percent loss of *ura4*<sup>+</sup> in diploids |
| --- | --- | --- |
| Strain | Treatment | % *ura4*Δ |
| Haploid | 0.0025% MMS | 0% (0/12) |
| Mat2-102 het. | No treatment | 87% (20/23) |
| Mat2-102 het. | 0.0025% MMS | 100% (22/22) |
| Proto. het. | No treatment | 96% (23/24) |
| Proto. het. | 0.0025% MMS | 100% (24/24) |

Percent of 5-FOA resistant colonies that had completely lost the *ura4*<sup>+</sup> marker resulting in loss of heterozygosity. Colonies that maintained heterozygosity of *ura4*<sup>+</sup> have presumably lost *ura4*<sup>+</sup> function by mutation.

## Notes

### Competing Interest Statement

The authors have declared no competing interest.

### Summary of Updates

Supplemental figures and table have been uploaded.

